# The PAS domain-containing protein HeuR regulates heme uptake in *Campylobacter jejuni*

**DOI:** 10.1101/057042

**Authors:** Jeremiah G. Johnson, Jennifer A. Gaddy, Victor J. DiRita

## Abstract

*Campylobacter jejuni* is a leading cause of bacterial-derived gastroenteritis. A previous mutant screen demonstrated that the heme uptake system (Chu) is required for full colonization of the chicken gastrointestinal tract. Subsequent work found identified a PAS domain-containing regulator, termed HeuR, as required for chicken colonization. Here we confirm that both the heme uptake system and HeuR are required for full chicken gastrointestinal tract colonization, with the *heuR* mutant being particularly affected during competition with wild-type *C. jejuni.* Transcriptomic analysis identified the chu genes-and those encoding other iron uptake systems-as likely regulatory targets of HeuR. Purified HeuR specifically bound the *chuZA* promoter region in electrophoretic mobility shift assays. Consistentwith a role forHeuR in *chu* expression, *heuR* mutants were unable to efficiently use heme asa source of iron in iron-limiting conditions and, mutants exhibited decreased levels of cell-associated ironby massspectrometry.Finally, we demonstrate that a heuR mutant of *C. jejuni* is resistant to hydrogen peroxide, and that this resistance correlates to elevated levels of catalase activity.

**Author Summary:** *Campylobacter jejuni* causes millions of gastrointestinal infection every year. This is primarily due to the its ability to reside in the gastrointestinal tract of chickens. *C.jejuni* contaminates chicken meat during harvesting and processing. Following consumption of undercooked chicken or uncooked food that was contaminated with raw chicken juice, humans develop a debilitating illness that is characterized by diarrhea and abdominal cramps. As chickens are the source of most human infections, there is a need to understand how *C. jejuni* colonizes chickens so we can develop ways to reduce its presence in chickens and thereby improve food safety. Most organisms require iron to thrive and that some bacteria steal iron from host molecules, including hemoglobin. Here we demonstrate that *C. jejuni* may need to get iron from hemoglobin in order to colonize the chicken and that aregulatory protein, HeuR, controls the ability ofthe bacteria to do this. If we can understand how this protein works, we may be able to develop ways to inhibit its function and reduce the ability of *C. jejuni* to get iron during chicken colonization. This would limit the amount of *C. jejuni* in the chicken and make food safer.

## Introduction

*Campylobacter jejuni* is a leading cause of gastrointestinal infection worldwide, with a projected 1.3 million annual cases in the United States (1).The prevalence of *C. jejuni* infection is due to its ability to reside asymptomatically within the gastrointestinal tract of avians, especially chickens. During processing of poultry, *C. jejuni* is released from the gastrointestinal tract, contaminating meat products as a result. Patients with *C. jejuni* infection typically develop mild to severe, bloody diarrhea that may be accompanied by fever. Additionally, several post-infectious disorders have been associated with *C. jejuni* infection, including Guillain-Barre Syndrome, post-infectious irritable bowel syndrome, and reactive arthritis (2, 3). Further highlighting these concerns is increasing resistance of *C. jejuni*to the clinically-relevant antibiotics,azithromycin and ciprofloxacin, prompting the Center for Disease Control and Prevention to list antibiotic-resistant *C. jejuni* as a“Serious Threat” to public health (1).

Due to the importance of chickens in human *C. jejuni* infection, much emphasis has been placed on identifying and characterizing factors that are required for colonization of the chicken gastrointestinal tract. Previously, our group has used both signature-tagged mutagenesis (STM) and insertion sequencing (TnSeq) approaches to identify colonization determinants of *C. jejuni* (4, 5). The earlier STM approach identified genes involved in chemotaxis and flagellar biogenesis, but several other colonization factors were identified using the TnSeq approach.One determinant identified to be reduced by 250-fold in cecal outputs by TnSeq was a component of the heme utilization gene cluster, *chuB* (Cjj81176_1602), indicating that acquiring iron from heme may be important during colonization of the chicken gastrointestinal tract (4).

Due to the prominent redox potential of Fe^2+^/Fe^3+^, most organisms require iron as a redox cofactor for enzyme complexes. Since the host often restricts iron availability to limit microbial growth-a process termed nutritional immunity-many bacterial species have developed methods for obtaining iron from host molecules, including heme (6). The heme utilization gene cluster of *C. jejuni*consists of genes encoding a predicted TonB-dependent heme receptor(*chuA*),a hemin ABC transporter permease (*chuB*),a hemin ABC transporter ATP-binding protein (chuC), and a periplasmic hemin-binding protein (*chuD*)(7). Divergently transcribed from the transport genes is a previously characterized heme oxygenase, *chuZ*, which directly binds heme and degrades it in the presence of an electron donor (7, 8). All components of the heme utilization gene cluster are required for efficient use of heme as a sole iron source and the ferric uptake repressor (Fur) protein directly binds to the promoter region of the gene cluster, implicating Fur as a *chu* locus regulator (7).

These data are consistent with observations that, in most organisms, intracellular iron levels are tightly regulated to avoid forming toxic hydroxyl radicals and superoxide anions via Haber-Weiss-Fenton catalysis (9). *C. jejuni* is exceptional in that it is also exquisitely sensitive to oxidative stress and, as a result, adequate intracellular iron levels must be maintained as a redox cofactor. While the above studies indicated that increased iron levels repress *chu* gene expression, likely via Fur binding, no positive regulators of heme and/or iron uptake have been identified. Due to the importance of iron in combating oxidative stress in *C. jejuni*, it would be surprising if *Campylobacter* did not have some mechanism for increasing iron uptake under unfavorable redox states. As such, we hypothesized that *C. jejuni* likely maintains intracellular iron levels using both negative regulation, as seen in previous work with Fur, and positive regulation.

In subsequent TnSeq experiments that aimed at finding additional chicken colonization determinants, a candidate regulator was identified, Cjj81176_1389 (unpublished data). This regulator is predicted to contain an N-terminal Per-Arnt-Sim (PAS) domain and a C-terminal helix-turn-helix DNA binding domain. The PAS domain is a ligand-binding domain that interacts with a chemically diverse array of small molecules (10). These domains, while highly divergent in primary sequence identity, maintain a conserved three-dimensional architecture (10). When ligand binds to a PAS domain, the protein either directly initiates a signaling response or the protein-ligand complex becomes capable of responding to a secondary cue, including gas molecules, photons, or redox potential (10). We reasoned that since the heme utilization gene cluster and Cjj81176_1389 appear to be required for full colonization of the chicken gastrointestinal, and that Cjj81176_1389 may be sensing intracellular redox potential, Cjj81176_1389 could be directly regulating *chu* gene expression.

Here we demonstrate that Cjj81176_1389 mutants are significantlyreduced for chicken gastrointestinal tract colonization, particularly during competition with wild-type *C. jejuni*. Subsequent transcriptomic analysis found that this regulator, hereafter referred to as HeuR (for heme uptake regulator), positively influences *chu* gene expression, as well as other iron uptake systems. Purified HeuR specifically binds the *chuZA* promoter region, indicating it may regulate *chu* gene expression directly. Further, *heuR* mutants are unable to efficiently use heme as a sole iron source and exhibit decreased levels of cell-associated iron. Finally, we show that loss of *heuR* leads to hydrogen peroxide resistance, in turn due to elevated levels of catalase activity.

## Results

***C. jejuni* lacking *heuR* are unable to efficiently colonize the chicken gastrointestinal tract.** Colonization studies using defined insertion-deletion mutants were performed using day-of-hatch chicks. Animals were inoculated with: 2.5×10^8^cfu wild-type *C. jejuni*, 1.7×10^8^ cfu *heuR* mutant, or 1.9×10^8^ *chuA* mutant. Following 7 days of colonization, viable bacteria present within cecal samples were enumerated and the medians found to be: 5.8×10^9^ cfu/g for wild-type *C. jejuni*, 6.6×10^8^ cfu/g for *heuR,* and 5.6×10^7^ cfu/g for the *chuA* mutant.The chicks that were mock infected with PBS did not yield detectable *C. jejuni*. Statistical analysis of the medians revealed that both *heuR* mutants were significantly decreased for colonization, but the *chuA* mutant was not (p = 0.1).

Competition analysis using the *heuR* mutant indicated asevere competitive defect of the mutant versus wild-type *C. jejuni*. The *heuR* mutant yielded a mean competitive index value of 0.004±0.007, representing a 250-fold decrease in colonization potential compared to wild-type *C. jejuni* (Figure 1B). Statistical analysis of this competitive disadvantage was found to be significant.

**Figure 1.**
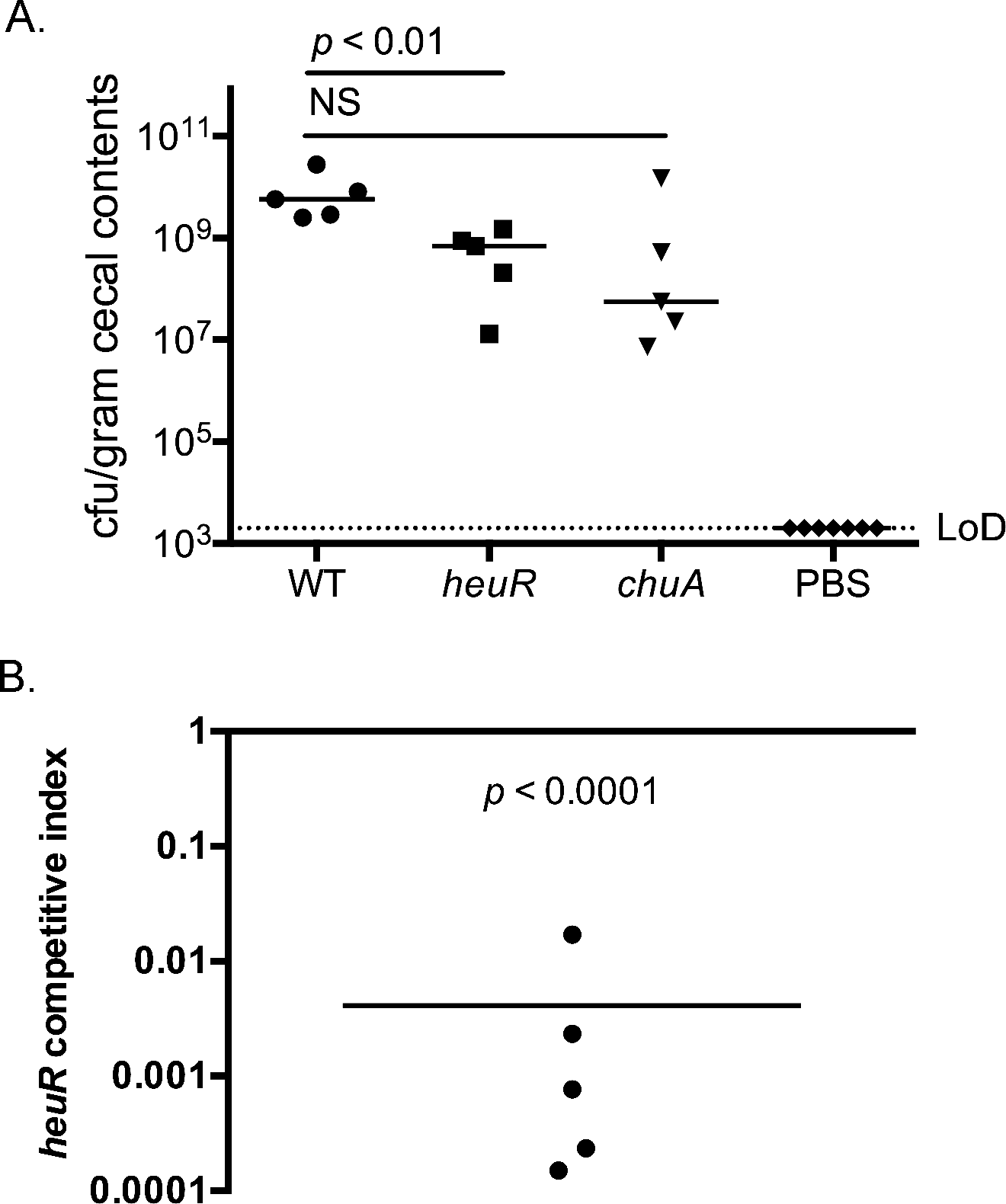
Colonization of day-of-hatch chicks with *heuR* mutants. (A)Mono-colonization of day-of-hatch white leghorn chicks with either wild-type *C.jejuni*, the *heuR* mutant,or the *chuA* mutant. Cecal loads were determined using selective media following a 7-day colonization. The median values are noted and results compared using the Mann-Whitney U-test. (B) Competition analysis of wild-type *C. jejuni* versus the *heuR*mutant. Following correction for inocula, the ratio of *heuR*/wild-type was determined and presented as a competitive index. Statistical analysis was performed using a one-sample t-test against a hypothetical value of 1.

**Genes involved in inorganic ion transport are underexpressed in the *heuR* mutant.** To determine the extent of the HeuR regulon, we employed transcriptomic analysis of wild-type *C. jejuni* and *heuR* mutant cultures. RNAseq analysis ofthe *heuR* mutant identified 43 genes whose transcripts were decreased in abundance by at least two-fold compared to wild-type *C. jejuni* (Table 1). These values represented statistically significant decreases for all genes identified. The Microbial Genomic context Viewer (MGcV) identified the NCBI GI numbers for 31/43 loci and, using a cluster of orthologous group (COG) analysis, assigned a functional class to 19/31 loci. Of the 19 loci for which a function could be assigned, eight (42%) were involved in inorganic ion transport and metabolism, including all five *chu* genes (Figure 2A). Also, amino acid transport and metabolism were well represented with four loci (21 %). The MGcV analysis did not identify NCBI GI numbers for the enterochelin system, which was also identified-iron acquisition by this system has been described in *Campylobacter* and multiple organisms (11, 12).

In addition to those genes that were found to exhibit reduced transcription in the *heuR* mutant, transcriptomic analysis identified 16 genes with elevated transcript abundance in the mutant (Table 2). The majority of this set included genes belonging to two gene clusters: Cjj81176_1390-1394 and Cjj81176_1214-1217. MGcV identified NCBI GI numbers for 11/16 genes and assigned functions to 10/11 loci. This group is fairly diverse with the largest representation belonging to the amino acid transport and metabolism COG (30%) (Figure 2B).

**Table 1.**
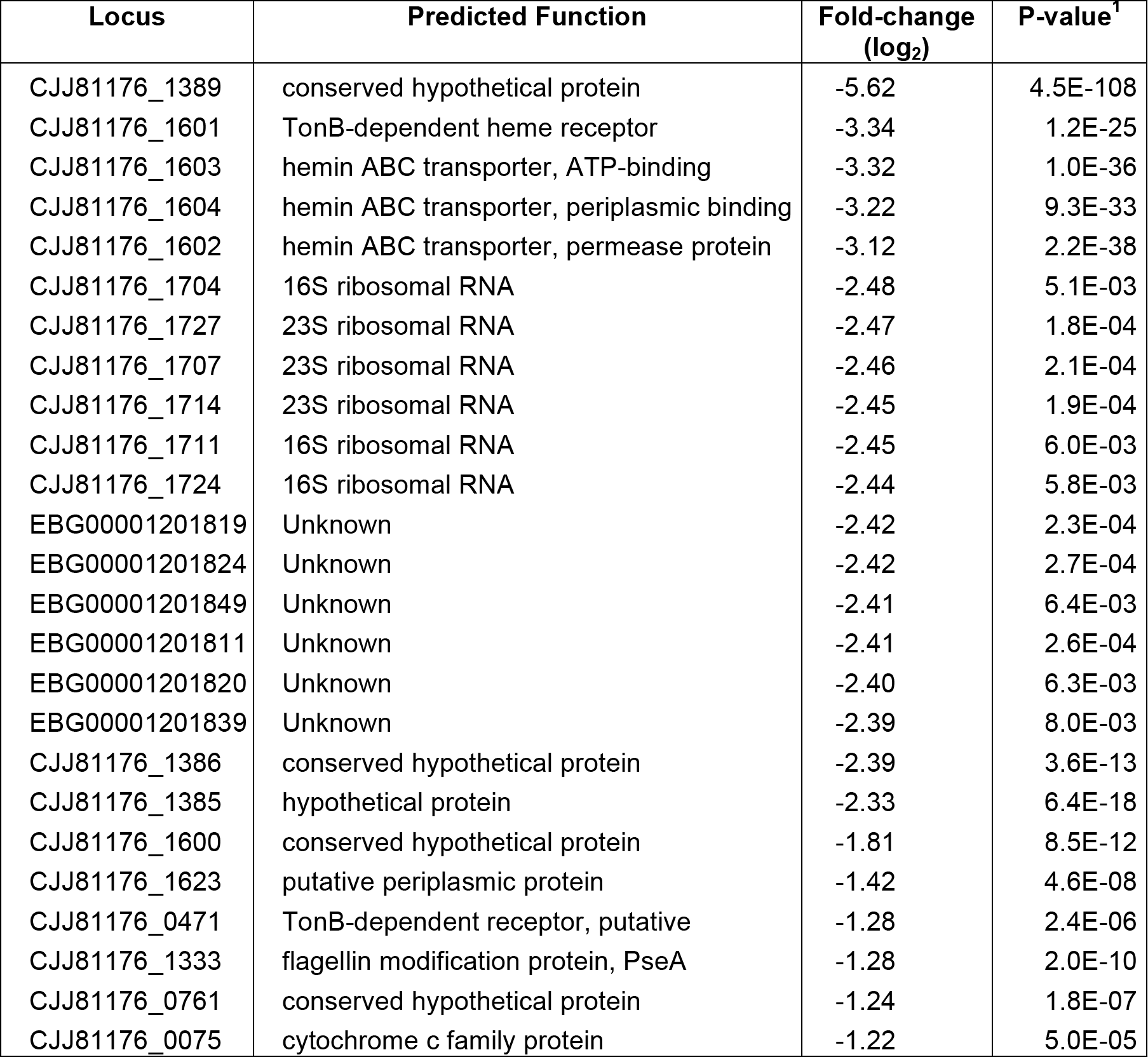
Genes with reduced transcript abundance in the *heuR* mutant.1.

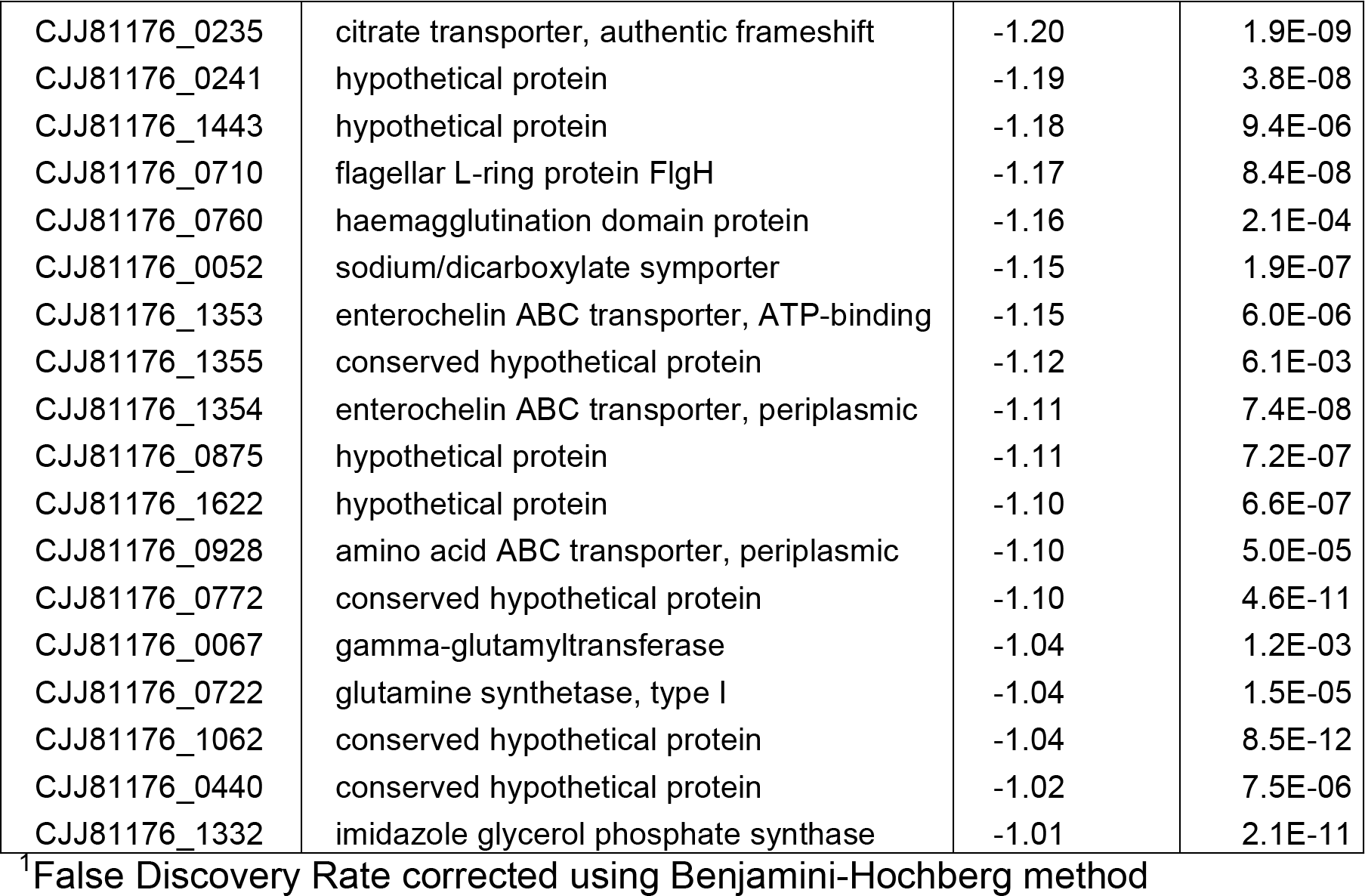

**Table 2.**
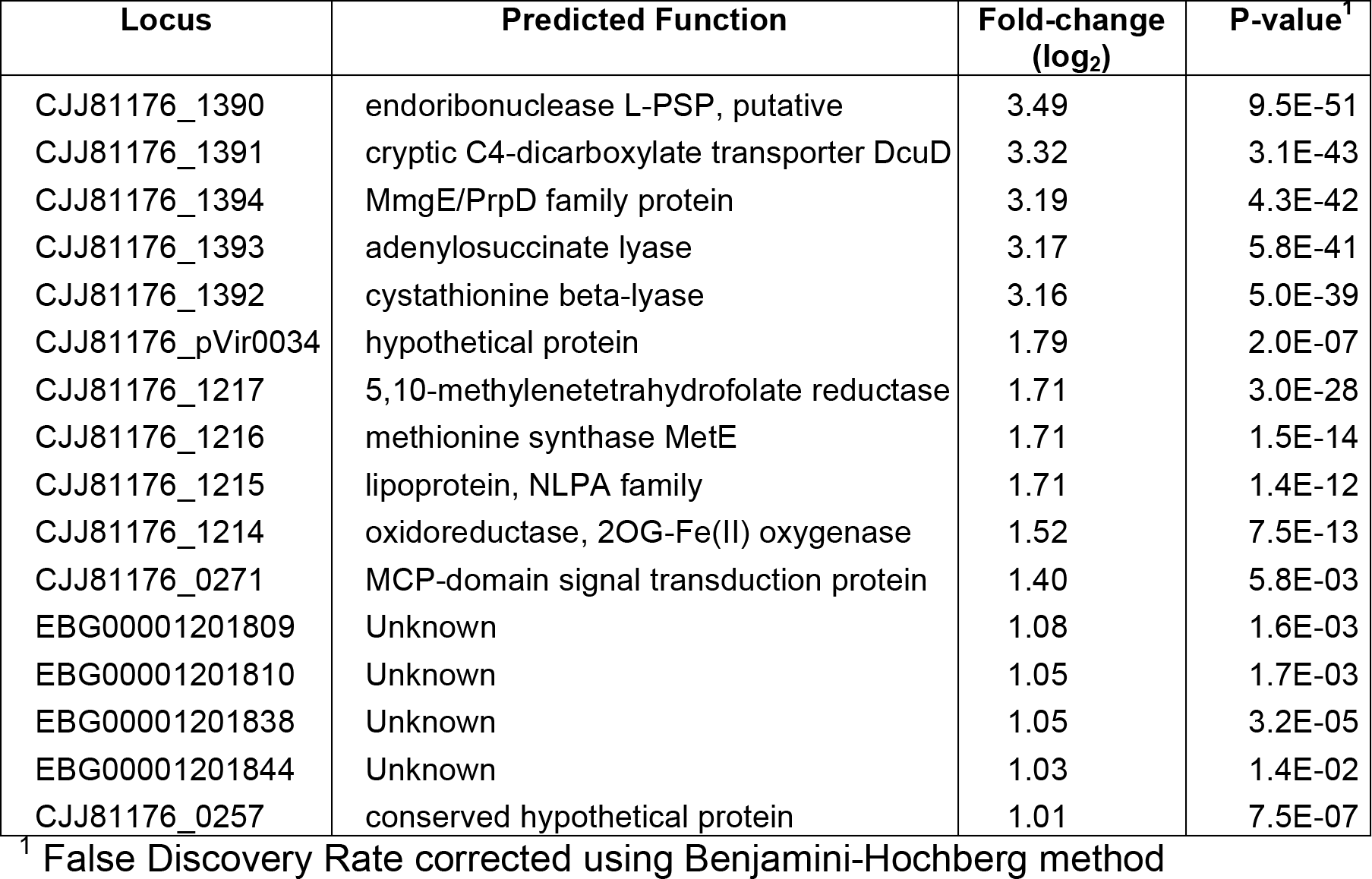
Genes with elevated transcript abundance in the *heuR* mutant.

| Locus | Predicted Function | Fold-change<br>(log<sub>2</sub>) | P-value<sup>†</sup> |
| --- | --- | --- | --- |
| CJJ81176_1390 | endoribonuclease L-PSP, putative | 3.49 | 9.5E-51 |
| CJJ81176_1391 | cryptic C4-dicarboxylate transporter DcuD | 3.32 | 3.1E-43 |
| CJJ81176_1394 | MmgE/PrpD family protein | 3.19 | 4.3E-42 |
| CJJ81176_1393 | adenylosuccinate lyase | 3.17 | 5.8E-41 |
| CJJ81176_1392 | cystathionine beta-lyase | 3.16 | 5.0E-39 |
| CJJ81176_pVir0034 | hypothetical protein | 1.79 | 2.0E-07 |
| CJJ81176_1217 | 5,10-methylenetetrahydrofolate reductase | 1.71 | 3.0E-28 |
| CJJ81176_1216 | methionine synthase MetE | 1.71 | 1.5E-14 |
| CJJ81176_1215 | lipoprotein, NLPA family | 1.71 | 1.4E-12 |
| CJJ81176_1214 | oxidoreductase, 2OG-Fe(II) oxygenase | 1.52 | 7.5E-13 |
| CJJ81176_0271 | MCP-domain signal transduction protein | 1.40 | 5.8E-03 |
| EBG00001201809 | Unknown | 1.08 | 1.6E-03 |
| EBG00001201810 | Unknown | 1.05 | 1.7E-03 |
| EBG00001201838 | Unknown | 1.05 | 3.2E-05 |
| EBG00001201844 | Unknown | 1.03 | 1.4E-02 |
| CJJ81176_0257 | conserved hypothetical protein | 1.01 | 7.5E-07 |

<sup>†</sup> False Discovery Rate corrected using Benjamini-Hochberg method

**Figure 2.**
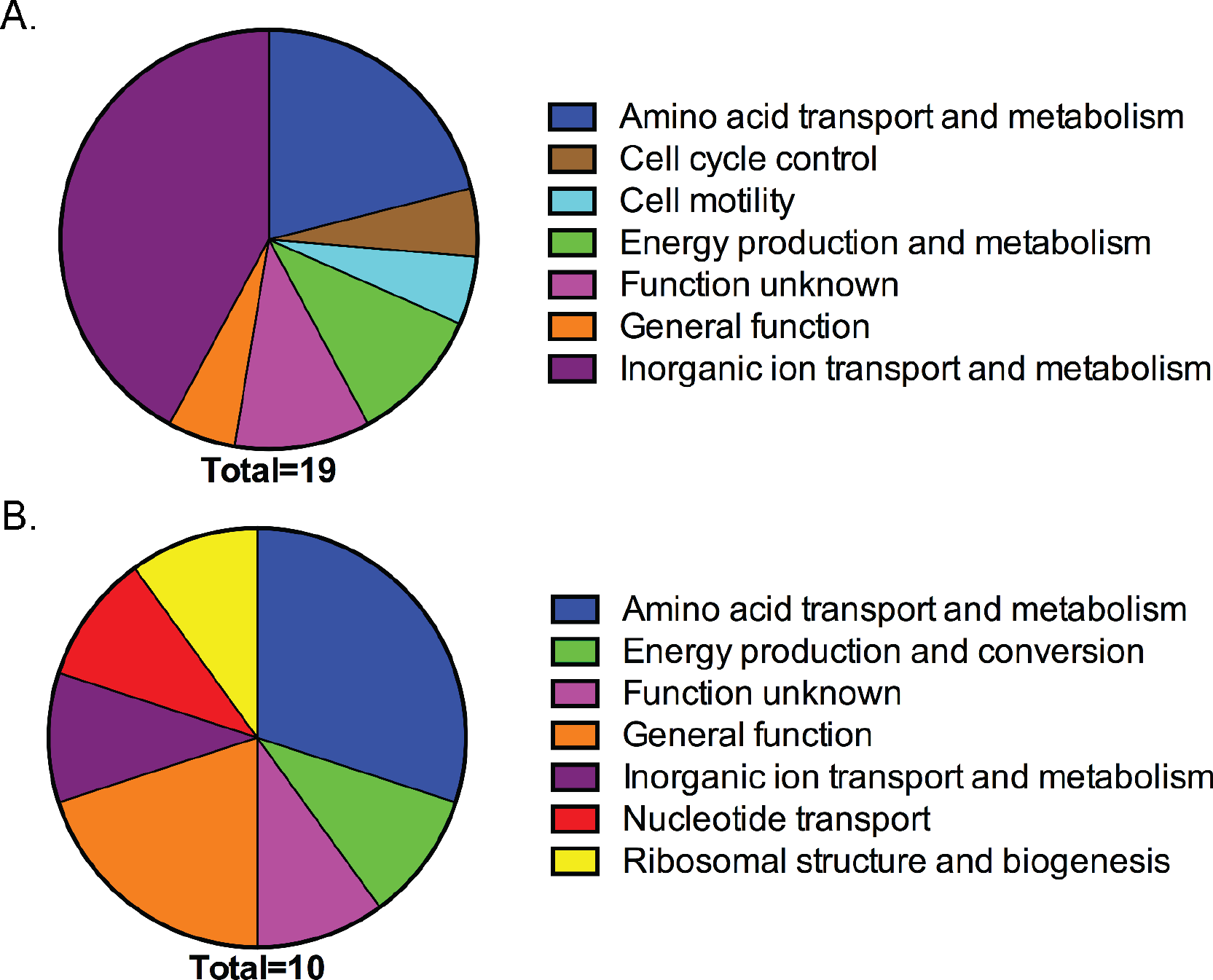
COG analysis of genes differentially expressed in the *heuR* mutant. (A) Identified COGs for genes in Table 1 that were found to be underexpressed in the *heuR* mutant. (B) COGs identified for genes in Table 2 that were overexpressed in the *heuR* mutant.

**Purified HeuR binds specifically to the *chuZA* promoter**. The putative structure of HeuR includes a C-terminal helix-turn-helix DNA binding domain. Given the transcriptome data demonstrating that *chu* RNA abundance is dependent on HeuR, we hypothesized that HeuR directly regulates *chu* gene expression. Purification of 6xHis-tagged HeuR was successful as soluble protein was obtained at concentrations of 4.3 mg/ml. Incubation of purified 6×His-HeuR with radiolabeled *chuZA* promoter and subsequent resolution by native polyacrylamide gel electrophoresis indicated binding of the protein to the *chuZA* probe using 30μg of 6×His-HeuR (Figure 3). The shifted species increased in intensity until 125μg of HeuR was used and the fragment representing unbound *chuZA* promoter was not detected. These results were in contrast to those obtained for a radiolabeled fragment that represented the intergenic region between *C. jejuni* 81-176 *mapA* and *ctsW,* which are not predicted to be regulated by HeuR. Mobility of this probe was relatively unaffected until 500μg of HeuR was used, when unbound probe intensity was modestly decreased.

**Figure 3.**
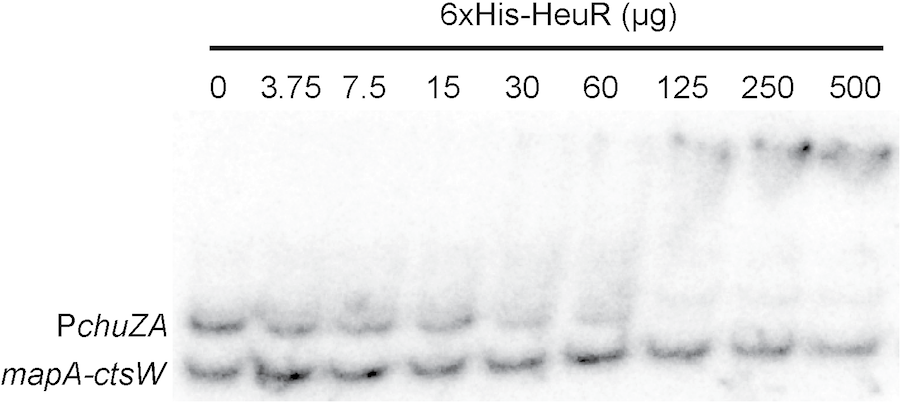
Binding of purified 6×His-HeuR to the *chuZA*promoter. Increasing amounts of purified 6×His-HeuR were added to binding reactions containing approximately 0.25nM of the *chuZA* promoter fragment and the *mapA-ctsW* intergenic region as a non-specific control.

**Mutants of *heuR* are unable to use heme as a source of iron**. To determine whether decreased expression of the Chu system affected the ability of the *heuR* mutant to use heme as an iron source, cultures of each strain were grown in iron-chelated conditions and supplemented with heme, hemoglobin, or ferric chloride. Both the *heuR* and *chuA* mutants were hypersensitive to iron-limiting conditions when compared to wild-type *C. jejuni* (Supplemental Figure 1). At 160μM desferrioxamine (DFOM), the *heuR* mutant with empty vector and the *chuA* mutant with empty vector exhibited greatly reduced growth compared to their behavior in iron-replete medium (Figure 4A). Heme added to these cultures was unable to rescue growth of the *heuR* and *chuA* mutants. Complementation of the *heuR* mutant with cloned *heuR* restored the ability to use heme as an iron source. In contrast, cloned *heuR* introduced into the *chuA* mutant did not restore heme dependent growth. Similar results were also seen using whole hemoglobin (Figure 4B). That iron restriction is the explanation for these phenotypes was supported by the restoration of growth in all strains through the addition of ferric chloride (Figure 4C).

**Figure 4.**
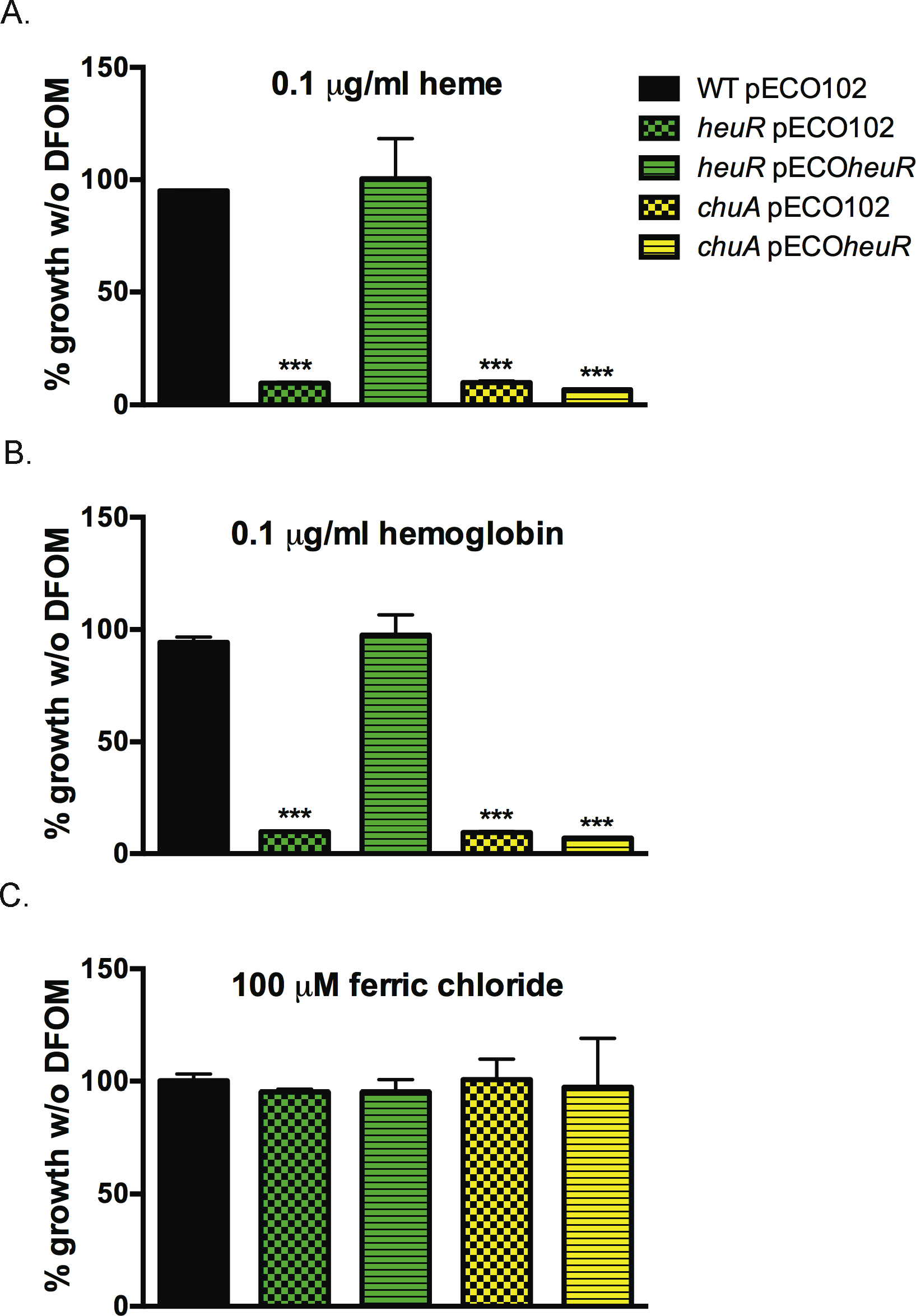
The *heuR* mutant is deficient in using heme as an iron source. (A) Growth of strains in media containing 160¼M desferrioxamine and 0.1 ¼g/ml purified heme. Represented as percent growth of same strains in media without desferrioxamine. (B) Growth of strains in media containing 160¼M desferrioxamine, but with 0.1¼g/ml purified hemoglobin. Represented as percent growth of strains in media without desferrioxamine. (C) Control growth of strains in media containing 160¼M desferrioxamine, but with 100¼M ferric chloride. Also represented as percent growth of strains in media without desferrioxamine. Statistical analysis performed using Student’s t-test;***: *p*<0.0001.

**Mutation of *heuR* results in decreased cell-associated iron**. The *heuR* mutant expressed lower abundance of mRNA from several iron uptake systems (Table 1), suggesting it may play a general role in regulating iron acquisition. To test this, we examined cell-associated iron levels using inductively coupled plasma mass spectrometry (ICP-MS) of wild-type *C. jejuni*, *heuR* mutant, and *chuA* mutant lysates. This analysis demonstrated that iron levels of the *heuR* and *chuA* mutants significantly decreased compared to those of wild-type *C. jejuni* (Figure 5A).

Iron levels in samples of the *heuR* mutant complemented with cloned *heuR* were similar to those of wild-type *C. jejuni* (Figure 5B) (p = 0.28). Similar to the results of the mutation alone, iron levels in the *chuA* mutant with pEC0102 were significantly decreased compared to wild-type *C. jejuni*. Consistent with our hypothesis that HeuR regulates iron acquisition through *chu* expression, when *heuR* was expressed in the *chuA* mutant, cell-associated iron remained significantly decreased compared to wild-type *C. jejuni* (Figure 5B).

**Figure 5.**
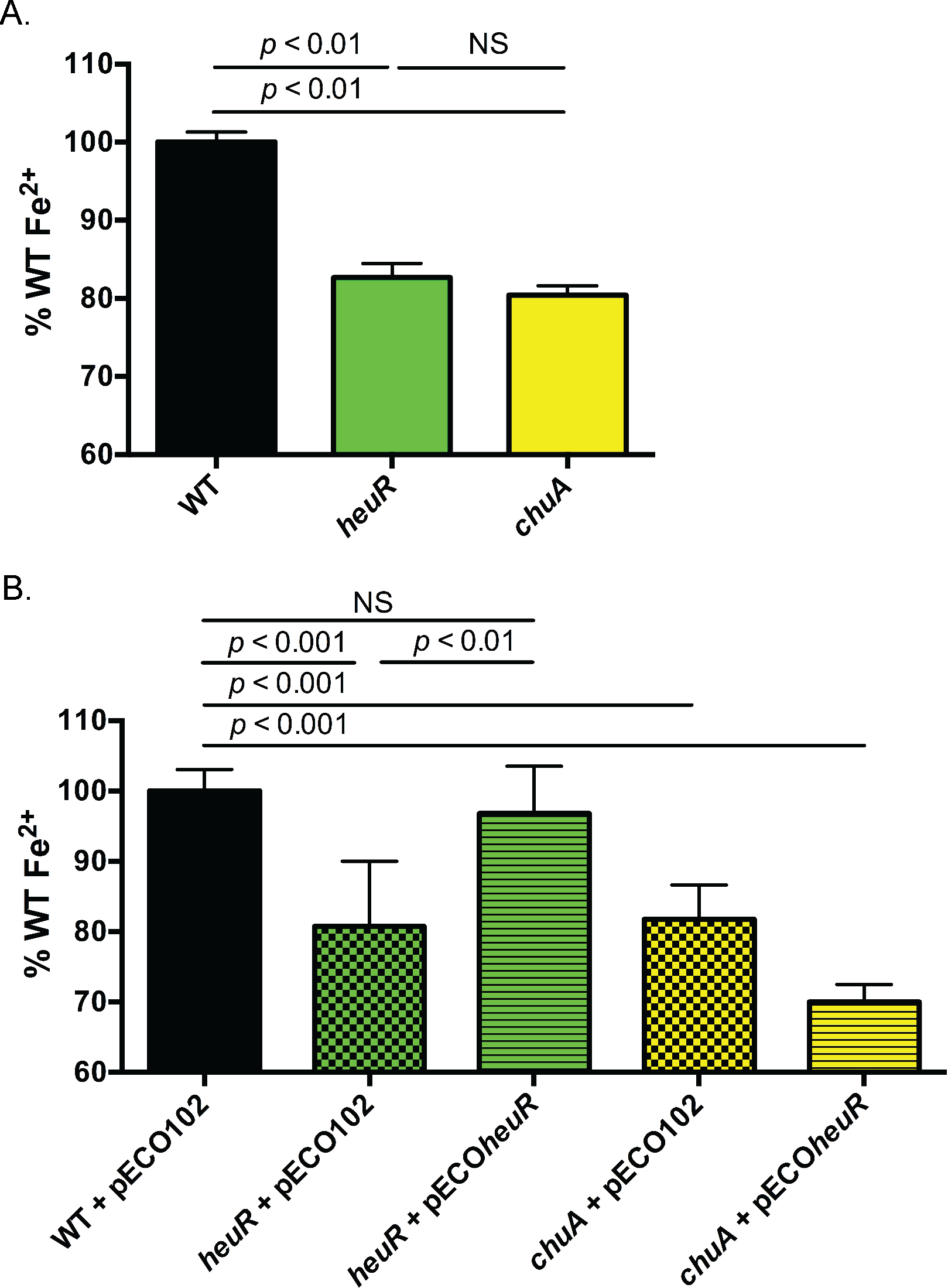
ICP-MS analysis of *C. jejuni* cell-associated iron. (A) Levels of cell-associated iron in wild-type *C. jejuni*, the *heuR* mutant, and the *chuA* mutant. Parts-per-billion iron levels for each strain were expressed asa percentage of the average iron levels observed in wild-type *C. jejuni*. (B) Complementation analysis of iron levels in strains carrying the empty vector control (pECO102) or the complementation construct (pECOheuR). Parts-per-billion iron levels for each strain were expressed as a percentage of the average iron levels observed in wild-type *C. jejuni* with empty vector. Statistical analyses performed using Student’s t-test.

**The *heuR* mutant exhibited increased resistance to hydrogen peroxide.** We observed a lack of alpha-hemolysis on blood plates on which the *heuR* mutant was grown (data not shown). As alpha hemolysis is due to the formation of methemoglobin in the presence of hydrogen peroxide, we hypothesized that the *heuR* mutant may degrade hydrogen peroxide more readily than wild-type *C. jejuni*. This was tested directly by incubating the *heuR* mutant in the presence of hydrogen peroxide, which revealed that the mutant is resistant. Growth of wild-type *C. jejuni* was generally limited by hydrogen peroxide at concentrations above 0.15mM and exhibited an IC50 value of 0.233mM (Figure 6A). In contrast, the *heuR* mutant was unaffected by hydrogen peroxide with significantly increased growth compared to wild-type *C. jejuni* at concentrations above 0.15mM. As a result, the IC50 value was not experimentally determined and by extrapolation was found to be 5.8mM. The *chuA* mutant exhibited an intermediate phenotype in the presence of hydrogen peroxide where growth remained relatively unaffected until concentrations reached 0.60mM, which was significantly more growth than that of wild-type *C. jejuni*. As such, the *chuA* mutant exhibited an IC50 of 0.526mM,a more than two-fold increase compared to wild-type *C. jejuni*, but approximately 10-fold lower than the *heuR* mutant.

Complementation of the *heuR* mutant with cloned *heuR* resulted in a slight hypersensitivity compared to wild-type*C. jejuni* with an IC50 value of 0.203mM (Figure 6B).

**Figure 6.**
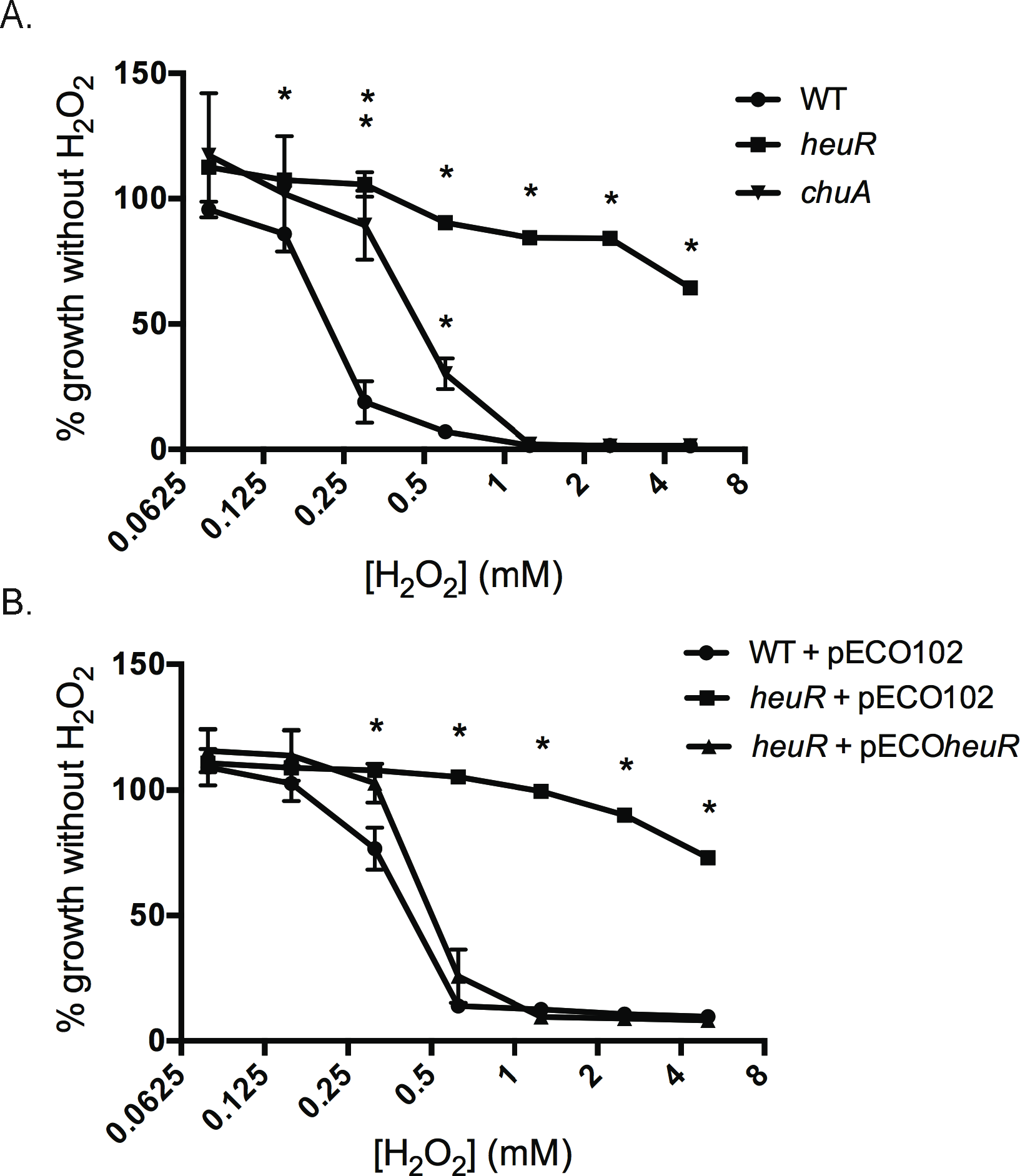
Dose-response analysis of *C. jejuni* mutants grown in presence of hydrogen peroxide. (A) Growth of wild-type *C. jejuni*, the *heuR* mutant, and the *chuA* mutant in response to increasing concentrations of hydrogen peroxide. (B) Complementation analysis of hydrogen peroxide resistance conferred by the *heuR* mutation. Wild-type with empty vector, the *heuR* mutant with empty vector, or the *heuR* mutant with cloned *heuR* were grown in the presence of increasing hydrogen peroxide concentrations. All growth is expressed as a percentage of that observed for each strain in media alone. Statistical analysis performed using a Student’s t test;*: *p*<0.01.

**Resistance of the *heuR* mutant to hydrogen peroxide is correlated with increased catalase activity**. We hypothesized that the elevated resistance of the *heuR* mutant to hydrogen peroxide may result from increased catalase activity. To determine whether this was the case, suspensions of strains were normalized and added to 30% H_2_O_2_, revealing that wild-type *C. jejuni* carrying empty vector was normally weakly positive (Table 3). In contrast, the *heuR* mutant transformed with empty vector was strongly positive for catalase activity. When *heuR* was re-introduced into the *heuR* background, catalase activity decreased and cells resembled wild-type activity. Also, similar to wild-type *C. jejuni*, the *chuA* mutant remained weakly positive, though the *chuA* mutant did appear to exhibit more catalase activity than wild-type *C. jejuni*.

**Table 3.** Catalase activity of *C. jejuni* strains.

| Strain (plasmid) | Catalase activity |
| --- | --- |
| Wild-type *C. jejuni* | +/- |
| *heuR* | ++++ |
| *chuA* | + |
| Wild-type *C. jejuni* (pECO102) | +/- |
| *heuR* (pECO102) | ++++ |
| *heuR* (pECO*heuR*) | +/- |

## Discussion

Using a TnSeq approach, we identified several determinants required by *C. jejuni* for efficient colonization of the chicken gastrointestinal tract (4). One determinant was a component of the *Campylobacter* heme utilization gene cluster, *chuB.* In this work, we describe a regulator, which we have named HeuR, which is also required for wild-type levels of colonization of the chicken gastrointestinal tract. Additionally, because HeuR is predicted to contain a PAS domain-a protein signature often associated with gas molecule and redox potential sensing-and that it and *chu* are both required for chicken colonization, we hypothesized that HeuR regulates heme uptake through *chu* gene expression.

Recently, HeuR was also found to control expression of Cjj81176_1390, as the gene product was overexpressed in a *heuR* mutant (13). The group that identified this phenotype named HeuR the *<u>C</u>ampylobacter* <u>f</u>lagella <u>i</u>nteraction <u>r</u>egulator (CfiR) due to flagellar interactions that occurred in response to gene disruption. This group further found that the gene product of Cjj81176_1390 forms dimers linked by a disulfide bridge that maybe responsive to the redox status of the cell (13). Our observations, recounted below, show that HeuR controls more genes than just those that promote flagellar interactions, including many that may be involved in combating oxidative stress (Figure 7). As such, we suggest that the designation CfiR does not accurately represent the full regulatory role in the bacterium and it is our recommendation that the name be changed to HeuR.

**Figure 7.**
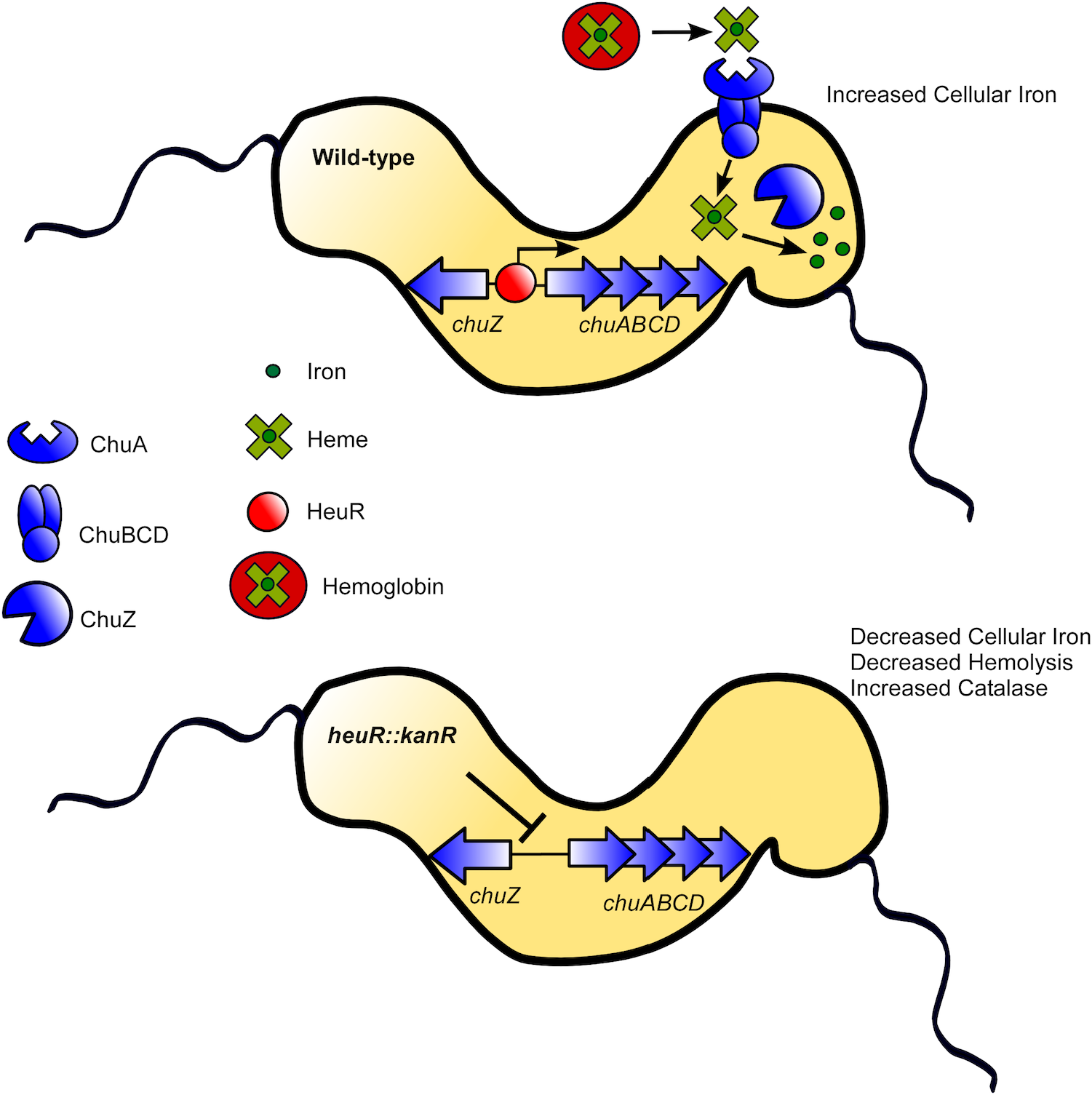
Model of HeuR regulation of the chuABCDZ loci and heme homeostasis in *C. jejuni.* Within the vertebrate host, heme is liberated from hemoglobin in an undefined manner. HeuR binds the intergenic region between chuZ and chuABCD to promote expression of heme acquisition functions. ChuA is the outermembrane heme receptor. ChuBCD transport heme to the cytoplasm of the cell where iron is liberated from bythe hemeoxygenase ChuZ. The *heuR* mutant exhibits decreased expression of the chuABCDZ loci, decreased ability to use heme and hemoglobin asa source of nutrient iron, and decreased cell-associated iron levels as aconsequence. Additionally, the *heuR* mutant exhibits decreased alpha hemolysis and increased catalase production.

As mentioned above, by RNAseq analysis of a *heuR* mutant strain, we identified several genes whose mRNA products were significantly less abundant in this background. Several of these were found to be involved in inorganic ion transport and metabolism, including all the genes in the previously identified *Campylobacter* heme utilization *(chu)* gene cluster. Transcript levels of several genes involved in enterochelin transport were decreased in the mutant strain, but were not identified by COG analysis as being involved in ion transport. As a result, this COG analysis is not comprehensive as the number of genes in the HeuR regulon that are involved in iron transport and metabolism is likely underreported. Nevertheless, based on these results, it becomes apparent that HeuR is primarily involved in maintaining sufficient levels of intracellular iron by upregulating systems that can scavenge for iron from diverse sources, which is supported by our ICP-MS data.

In addition to those genes that were underexpressed, transcripts of several were more abundant in the *heuR* mutant. Most of these genes are in two gene clusters: Cjj81176_1390-1394 and Cjj81176_1214-1217. The former, Cjj81176_1390-1394, includes the overexpressed gene product (Cjj81176_1390) noted above that promotes flagellar interactions, supporting both the previous study and our work (13). The latter gene cluster that is upregulated includes one gene that encodes an additional ion transporter and multiple genes predicted to be involved in methionine biosynthesis. The methionine biosynthetic enzymes encoded by these genes have been shown in other organisms to be sensitive to oxidation/oxidative stress (14). In an oxidative environment, these enzymes may need to be replenished, and their inverse expression with that of iron scavenging systems may be common. Whether upregulation of these genes is adirect or indirect effect of HeuR is currently unknown.

Standard growth assays using the iron chelator, desferrioxamine, found that both the *heuR* and *chuA* mutants are reduced for growth in iron-chelated conditions. As this was observed in media without the addition of hemoglobin, it may indicate that the Chu system is also partially responsible for the transport of free iron into the cell. In this case, it would likely be an accessory system since growth was completely restored in these mutants when iron was supplemented back, indicating the presence of a robust, primary iron transport system. When heme and hemoglobin were added to these mutants, growth was not restored, supporting the conclusion that both are required for using heme as an iron source. Re-introduction of *heuR* resulted in the ability to use heme as an iron source, likely through the Chu system, as overexpressing*heuR* in the *chuA* mutant did not enable the strain to use hemoglobin.

Analysis of cell-associated levels of iron showed that the *heuR* mutant is significantly decreased for iron and that these levels resemble those of a *chuA* mutant. Most of the iron reduction observed in these experiments is likely due to decreased heme utilization, as cultures were obtained from media containing sheep’s blood. This is further supported by the observation that complementation with *heuR* restores iron levels in a *heuR* mutant background, but was unable to do so in a *chuA* mutant background. As our data indicates a minor role for HeuR and the Chu system in acquiring iron independently of heme, we are currently determining whether decreased iron is apparent in the *heuR* and *chuA* mutants grown in media without heme or hemoglobin. This will help us determine to what extent other iron sources are impacting cellular iron levels in these mutants.

When the *heuR* mutant was grown on blood plates, differences in the hemolytic pattern relative to wild-type *C. jejuni* became apparent. Typically, *C. jejuni* exhibits alpha hemolysis on blood plates, likely due to the production of hydrogen peroxide that oxidizes the hemoglobin in the blood to methemoglobin (15). Alpha hemolysis was not observed in the*heuR* mutant, indicating that this mutant either produces less hydrogen peroxide or exhibits increased catalase activity. The mutant was highly resistant to hydrogen peroxide, which was associated with elevated levels of catalase activity; re-introduction of *heuR* led to both decreased resistance to hydrogen peroxide and catalase activity. Of the genes known to regulate catalase activity in *C. jejuni*-including *ahpC, fur, katA* and *perR*-none were found to be significantly up-or down-regulated in the *heuR* mutant (data not shown). Also, since iron levels are similar in the *chuA* mutant, it is unlikely that iron levels are responsible for the elevated catalase activity. Thus we hypothesize that HeuR regulates the production of an unknown factor that perhaps post-transcriptionally regulates the activity of *C. jejuni* catalase. We are currently working to determine what this factor is.

The *heuR* mutant exhibited significant decreases in colonization of the chicken cecum. This leads to two questions: i) is this system mainly required for combating oxidative stress in the cecum or is it used for other cellular processes and ii) is the system acquiring iron from a heme source or does it serve the dual purpose of acquiring free iron from the environment.

Oxidative stress in the cecum may come from a variety of sources, like metabolism by the intestinal microbiota or from the generation of reactive oxygen species due to intestinal inflammation. While the *Campylobacter-chicken* relationship is often considered to be commensal in nature, *C. jejuni* colonization of the chicken gastrointestinal tract may result in low levels of transient inflammation (Davis and DiRita, unpublished) (16–18). Also, while not mutually exclusive, iron acquisition systems like Chu may be more important for acquiring iron that can be used in other cellular processes including energy generation and DNA replication (6). We are currently working to determine which of the processes during chicken colonization iron is most important for.

Second, the question arises as to whether *C. jejuni* uses the Chu system to acquire iron from either/or a heme or non-heme source. It is currently unknown how much heme chickens consume in a commercial diet and we could find no data on heme availability in their natural diet. With the exception of higher plants that can produce heme in the plastids,the absence of heme molecules in most grains, plants, and insects make it less obvious how chickens would have access to heme-containing molecules (19). The likelihood then becomes that *C. jejuni* uses this system to either scavenges heme from a microbial source or from the host. The *C. jejuni* 81-176 genome encodes all heme biosynthetic enzymes with the exception of homologs for protoporphyrinogen oxidase (*hemY* or *hemG*). As such, in an iron rich environment, *C. jejuni* can likely synthesize its own heme. It could then use the heme uptake system to recover heme from other *Campylobacter* or from heme producing residents of the chicken microbiota. Lastly, it remains possible that this system is able to transport free iron. While the heme receptor and heme oxygenase (ChuA and ChuZ, respectively) would likely not be needed for free iron transport, the accompanying ABC transport system (ChuBCD) may be promiscuous and function similar to other iron transporters. Weare currently investigating whether the ChuBCD transporter can translocate non-heme iron into *C. jejuni*.

## Materials and Methods

**Day-of-hatch chick colonization.** Day-of-hatch white leghorn chickens were inoculated by oral gavage with 100¼l of an approximately 1×10^9^ cfu/ml bacterial suspension in sterile PBS (10^8^ cfu). Colonized chickens were housed for 7 days, euthanized, and cecal contents were collected. Cecal contents were homogenized in sterile PBS and serial dilutions were plated on *Campylobacter* selective media: MH agar containing 10% sheep’s blood, cefoperazone (40 μg/ml), cycloheximide (100 μg/ml), trimethoprim (10 μg/ml), and vancomycin (100 μg/ml). Plates were grown for 48h under microaerobic conditions and colonies enumerated. Statistical analysis of the medians was performed using a Mann-Whitney test.

For the competition experiment, equal numbers of wild-type *C. jejuni* and the *heuR* mutant were mixed and approximately 1×10^8^ of each were used to inoculate day-of-hatch chicks by oral gavage. Harvest of cecal contents was done the same as above except serial dilutions were plated on *Campylobacter* selective media with and without kanamycin (100 μg/ml). Competitive index values were calculated following correction for inoculum amounts by dividing the number of *heuR* mutants by the number of wild-type bacteria. Statistical analysis was done using a one-sample t-test against a hypothetical value of 1.

All chicken experiments and protocols were approved by the Institutional Animal Care and Use Committee in the Animal Care Program at Michigan State University.

**RNAseq analysis**. Triplicate cultures of wild-type DRH212 and the *heuR* mutant were grown on MH agar supplemented with 10% whole sheep’s blood and 10 μg/ml trimethoprim overnight at 37°C under microaerobic conditions. Cells were harvested and grown on new media for a second night using the same conditions. Following growth, cells were harvested from each individual plate and RNA was extracted using a previously described protocol (20). DNA was degraded using Turbo DNA-free (Thermo) according to the manufacturer’s instructions and removal was confirmed by conventional PCR. RNA quality was assessed using an Agilent 2100 Bioanalyzer and libraries were generated using the TruSeq RNA Library Preparation Kit (Illumina). Libraries were sequenced using an Illumina HiSeq 2000 formulated for single-end, 50-nucleotide reads.

Analysis of sequence data was performed using SPARTA: Simple Program for Automated reference-based bacterial RNAseq Transcriptome Analysis (21). Briefly, raw reads were trimmed and assessed for quality using Trimmomatic and FastQC, respectively (22). Trimmed reads were mapped to the *C. jejuni* 81-176 reference genome using Bowtie and expression levels determined by HTSeq (23, 24). Lastly, differential expression analysis was performed using edgeR (25). Cluster of orthologous group (COG) identification of selected genes was performed using Microbial Genomic context Viewer (MGcV).

**Protein purification**. The gene *heuR* was amplified by PCR from *C. jejuni* 81-176 genomic DNA using primers 5’Cjj1389 *BamHI* and 3’Cjj1389 *SalI.* The PCR fragment was subcloned into pGEM-T Easy using the manufacturer’s protocol and sequenced using primers T7 and SP6. Fragments that were confirmed to have maintained the full nucleotide sequence of *heuR* were excised using *BamHI* and *SalI* and ligated into the expression vector, pQE-30 (Qiagen). Ligation products were transformed into *E. coli* NEB C3013 and insertion was confirmed using restriction digest of purified plasmids.

Induction of 6×His-HeuR using 1 mM IPTG was confirmed for six constructs by SDS-PAGE and Coomassie staining. One strain was used for protein purification where a 1L culture was grown to mid-log and induced for 3hat 37°C using 1mM IPTG. Cells were pelleted at 8,000 rpm for 20 minat 4°C and re-suspended in 20 ml lysis buffer (pH 8.0) (50 mM NaH_2_PO_4_, 300 mM NaCl, 10 mM imidazole). The cell suspension was passed three times through a french press and cell debris was removed by centrifugation at 8,000 rpm for 20 min at 4°C. The cleared lysate was added to 5 ml of NiNTA agarose that was preequilibrated with lysis buffer and rocked overnight at 4°C. This resin-protein mixture was packed by gravity into a chromatography column and lysate was passed through and collected. Non-specific proteins were removed by washing the resin bed twice with 5 column volumes (25 ml) of wash buffer (pH 8.0) (50 mM NaH2PO4, 300 mM NaCl, 20 mM imidazole). Bound protein was eluted four times using 1 ml of elution buffer (pH 8.0) (50 mM NaH_2_PO_4_, 300 mM NaCl, 250 mM imidazole). Successful purification of 6×His-HeuR was verified by SDS-PAGE and Coomassie staining. Fractions containing the highest amount of HeuR were pooled and dialyzed against storage buffer (50 mM Tris (pH 7.5), 150 mM NaCl, 0.5 mM EDTA, 0.02% Triton ×-100, 2 mM DTT, 50% glycerol). Concentrations of 6×His-HeuR were determined using a BCA Protein Assay (Pierce) and aliquots of protein were stored at-80°C.

**Electrophoretic mobility shift assays**. A specific DNA probe was generated from the *chuZA* promoter region and a non-specific probe was generated from the *mapA-ctsW* intergenic region by PCR of *C. jejuni* 81-176 genomic DNA. Amplicons were purified using a PCR Purification Kit (Qiagen) and subsequently end-labeled with [y-^32^P] ATP using T4 polynucleotide kinase (PNK) (New England Biolabs). DNA binding reactions containing radiolabeled specific (*chuZA* promoter) and non-specific probes (*mapA-ctsW* intergenic) at approximately 0.25 nM, DNA binding buffer (10 mM Tris, pH 7.5, 50 mM KCl, 1 mM EDTA, 1 mM DTT, 5% glycerol), 25 ng/μl poly(dA-dT), and 100 μg/ml BSA were incubated for 5 min at room temperature. Purified 6×His-HeuR was diluted two-fold in binding buffer and 1 μl containing 500, 250, 125, 60, 30, 15, 7.5, or 3.75 μg of protein was added to individual 19 μl binding reactions (20 μl total). These reactions-and a control reaction without 6×His-HeuR-were incubated for 15 min at room temperature and subsequently resolved on a 5% polyacrylamide native Tris-glycine gel at 4°C. Gels were imaged using aTyphoon phosphoimager.

**Hemoglobin and heme growth assays**. For bacterial growth assays, *C. jejuni* strains were grown overnight on tryptic soy agar plates supplemented with 5% sheep blood and appropriate antibiotics (chloramphenicol and/or kanamycin). The following day, plate cultures were utilized to inoculate 5 mL of Mueller-Hinton broth supplemented with appropriate antibiotics. These liquid cultures were incubated under shaking conditions at 37°C overnight. The following day, cultures were diluted 1:10 in fresh Mueller-Hinton broth (26) supplemented with appropriate antibiotics (chloramphenicol and/or kanamycin) alone (Medium Alone) or supplemented with 40, 80, 160¼M of the iron chelator desferrioxamine mesylate (Sigma Aldrich) alone or in combination with 0.1¼g/mL of purified human hemoglobin (1) or 0.1¼g/mL of hemin at 37°C in room air supplemented with 5% CO_2_. Bacterial growth was measured using a spectrophotometric reading at 600 nm (OD_600_). Statistical analysis was performed using a Student’s t-test.

**ICP-MS analysis**. Cultures of wild-type *C. jejuni* DRH212, the *heuR* mutant, and a *chuA* mutant, were grown overnight at 37°C under microaerobic conditions on MH agar with 10% whole sheep’s blood containing either 10 μg/ml trimethoprim or 100 μg/ml kanamycin-for mutant strains. For complementation analysis, wild-type with pECO102, the *heuR* mutant with pECO102, the *heuR* mutant with pECOheuR, the *chuA* mutant with pECO102, and the *chuA* mutant with pECO*heuR* were grown on MH agar with 10% sheep’s blood containing either chloramphenicol (30 μg/ml) alone-for wild-type-or chloramphenicol and kanamycin (100 μg/ml)-for the plasmid-containing mutants. Cultures were grown overnight under microaerobic conditions at 37°C. Strains were passed for a second night on fresh media and harvested for iron quantification. Approximately 10^10^ cells were washed three times with 5 ml and re-suspended in 500 μl of molecular-grade, distilled water. Cells were solubilized overnight at 50°C in 1ml 50% nitric acid, followed by the addition of 9 ml of water.

Extracts were analyzed for ^57^Fe using A Thermo Scientific ICAP Q quadrupole inductively coupled plasma mass spectrometer in kinetic energy dispersion mode. Instrument calibration was done with aliquots of a Fe synthetic standard solution diluted at 2.5, 10, and 100 ppb and natural water standards NIST SRM 1640 and 1643e. A solution of 100 ppb Fe was analyzed every 5-6 samples to monitor instrumental drift; all samples were corrected for linear drift. Oxides formation rate was determined to be<2% by measuring CeO/Ce and double-charged cation formation rate was found to be<3% by measuring 137Ba^++^/137Ba from a solution with approximately 1 ppb Ce and 1 ppb Ba.

Parts-per-billion levels for each strain were normalized to those observed for the wild-type equivalent in each experiment and the standard deviation was plotted. Statistical analysis was performed using a Student’s t-test.

**Hydrogen peroxide dose-response assays.** Cultures of wild-type *C. jejuni*, the *heuR* mutant, and the *chuA* mutant were grown overnight at 37°C under microaerobic conditions on MH agar with 10% sheep’s blood. Typically, *C. jejuni* exhibits alpha hemolysis on blood plates, likely due to the production of hydrogen peroxide that oxidizes the hemoglobin in the blood to methemoglobin. Strains were passed for a second night on new media under the same conditions. Cells were harvested into MH broth and normalized to an OD A_600_ of 0.05. These suspensions were added to equal volumes (100 pl:100 pl) of 2-fold dilutions of MH broth with hydrogen peroxide in a 96-well plate. Hydrogen peroxide concentrations ranged from 10 mM to 0.16 mM. This dilution resulted in bacterial concentrations of 0.025 OD and hydrogen peroxide concentrations ranging from 5 mM to 0.08 mM. These cultures were grown statically for 48h at 37°C under microaerobic conditions followed by re-suspension of bacteria and recording of terminal OD A_600_ values. IC50 values were calculated using non-linear regression analysis of normalized responses with variable slopes in GraphPad Prism 6. Statistical analysis of differences between strains at each concentration of hydrogen peroxide was performed using a Student’s t-test.

For complementation analysis of the *heuR* phenotype, the complemented strains used for ICP-MS analysis were subjected to the same hydrogen peroxide dose-response assay as those strains above. Statistical analysis was also performed using a Student’s t-test.

**Catalase assays.** Cultures of wild-type *C. jejuni*, the *heuR* mutant, and the *chuA* mutant were grown overnight at 37°C on MH agar with 10% sheep’s blood under microaerobic conditions. These strains were passed for a second night on new media under the same conditions and cells were harvested into sterile PBS. Care was given to avoid contaminating the suspensions with media since blood exhibits catalase activity. Suspensions were normalized to an OD A_600_ of 1.0 and 10 μl of suspension was added to 10 μl of 30% hydrogen peroxide (J.T. Baker) on a glass slide.Reactions were performed in triplicate and qualitatively noted.

## Acknowledgements

We would like to thank Ben K. Johnson at Michigan State University for his help with analyzing RNAseq results using SPARTA.1. JGJ was supported by USDA NIFA grant 2013-67012-25269. Funding for JAG was provided by Department of Veteran Affairs’ Office of Medical Research¼rant CDA-21K2BX001701. VJD was supported by start-up funds from Michigan State University and by NIH NIAID grant R21AI111192.

## Supplementary Figure

**S1.**
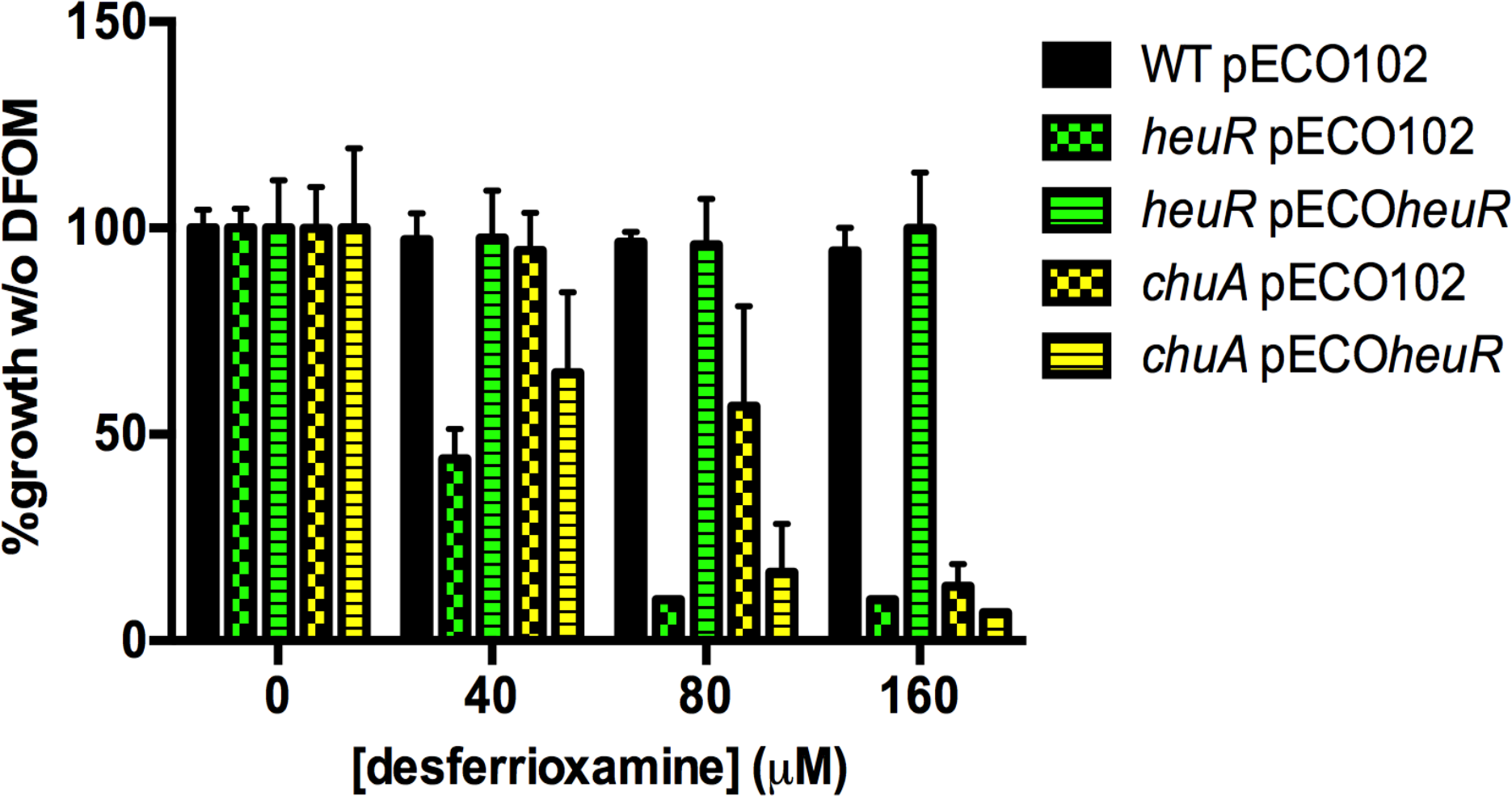
Dose-response analysis of desferrioxamine. *C. jejuni* strains grown in the presence of increasing amounts of desferrioxamine (DFOM). Growth at each concentration of DFOM is expressed as percent growth of that strain in media alone.

